# Developmental effects of maternal smoking during pregnancy on the human frontal cortex transcriptome

**DOI:** 10.1101/236968

**Authors:** Stephen A. Semick, Leonardo Collado-Torres, Christina A. Markunas, Joo Heon Shin, Amy Deep-Soboslay, Ran Tao, Laura J. Bierut, Brion S. Maher, Eric O. Johnson, Thomas M. Hyde, Daniel R. Weinberger, Dana B. Hancock, Joel E. Kleinman, Andrew E. Jaffe

**Affiliations:** Lieber Institute for Brain Development, Johns Hopkins Medical Campus, Baltimore, MD, 21205, USA; Center for Computational Biology, Johns Hopkins University, Baltimore, MD, 21205, USA; Behavioral and Urban Health Program, Behavioral Health and Criminal Justice Division, RTI International, Research Triangle Park, NC, 27709, USA; Department of Psychiatry, Washington University School of Medicine, St Louis, MO 63110, USA; Department of Mental Health, Johns Hopkins Bloomberg School of Public Health, Baltimore, MD, 21205, USA; Fellow Program and Behavioral Health and Criminal Justice Division, RTI International, Research Triangle Park, NC, 27709, USA; Department of Psychiatry and Behavioral Sciences, Johns Hopkins School of Medicine, Baltimore, MD 21205, USA; Department of Neurology, Johns Hopkins School of Medicine, Baltimore, MD, 21205, USA; Department of Neuroscience, Johns Hopkins School of Medicine, Baltimore, MD, 21205, USA; McKusick-Nathans Institute of Genetic Medicine, Johns Hopkins School of Medicine, Baltimore, MD 21205, USA; Department of Biostatistics, Johns Hopkins Bloomberg School of Public Health, Baltimore, MD, 21205, USA

## Abstract

Cigarette smoking during pregnancy is a major public health concern. While there are well-described consequences in early child development, there is very little known about the effects of maternal smoking on human cortical biology during prenatal life. We therefore performed a genome-wide differential gene expression analysis using RNA sequencing (RNA-seq) on prenatal (N=33; 16 smoking-exposed) as well as adult (N=207; 57 active smokers) human post-mortem prefrontal cortices. Smoking exposure during the prenatal period was directly associated with differential expression of 14 genes; in contrast, during adulthood, despite a much larger sample size, only 2 genes showed significant differential expression (FDR<10%). Moreover, 1,315 genes showed significantly different exposure effects between maternal smoking during pregnancy and direct exposure in adulthood (FDR<10%) – these differences were largely driven by prenatal differences that were enriched for pathways previously implicated in addiction and synaptic function. Furthermore, prenatal and age-dependent differentially expressed genes were enriched for genes implicated in non-syndromic autism spectrum disorder (ASD) and were differentially expressed as a set between patients with ASD and controls in post-mortem cortical regions. These results underscore the enhanced sensitivity to the biological effect of smoking exposure in the developing brain and offer novel insight into the effects of maternal smoking during pregnancy on the prenatal human brain. They also begin to address the relationship between *in utero* exposure to smoking and the heightened risks for the subsequent development of neuropsychiatric disorders.

**One Sentence Summary:** Maternal smoking during pregnancy alters the expression of genes within the developing human cortex and these changes are enriched for genes implicated in neuropsychiatric disorders.

## Introduction

Cigarette smoking and nicotine addiction continue to be major public health problems, due to their established association with increased risk of cancer, respiratory disease, and many other disease outcomes in adults (*1*). Cigarette smoking exposure also is highly deleterious to fetal development, having been associated with heightened risk of intrauterine growth restriction and low birth weight (*2*). There is accumulating evidence that *in utero* exposure to nicotine and other elements of cigarette smoking is associated with a greater risk of the subsequent development of mental illnesses (*3, 4*). Despite the well-known adverse effects of smoking on the smoker and the fetus carried by mothers who smoke, little is known about the effects of smoking on gene expression in the adult and especially the fetal brain.

In the United States, 8.4% of pregnant women smoke while pregnant (*5*). Smoking during pregnancy has been repeatedly associated with a host of adverse health outcomes for the infant including premature birth, low birth weight, stillbirth, and mortality (*6*). Maternal smoking during pregnancy poses a potent danger to the developing human brain because many components of cigarette smoke, including nicotine, have neuroteratogenic effects (*7*). There is mixed evidence for maternal smoking during pregnancy as a risk factor for neuropsychiatric disorders including schizophrenia (*3, 4*), attention-deficit / hyperactivity disorder (*8–12*), Tourette syndrome (*12, 13*), impaired cognitive development (*14, 15*), obsessive-compulsive disorder (*12*), and autism spectrum disorder (*16*).

The molecular mechanisms underlying risk for neuropsychiatric disease from maternal smoking during pregnancy remains elusive. Previous studies have found epigenetic changes in cord blood (*17, 18*) and differential gene expression in placental tissue (*19*). However, only one prior study has directly examined associations of maternal smoking during pregnancy with epigenetic changes in the prenatal human brain (*20*), and no study, to our knowledge, has characterized transcriptional changes in the prenatal brain. Characterizing the gene regulatory changes associated with smoking exposure in prenatal life may identify potential mechanisms related to disease risks later in life especially with regard to neuropsychiatric disorders. We can better understand the link between maternal smoking during pregnancy and future behavioral and cognitive sequelae by directly studying molecular changes in prenatal human cortical tissue exposed to cigarette smoke.

To characterize molecular changes in the developing prenatal brain associated with *in utero* smoking exposure, we analyzed RNA-seq data from post-mortem fetal human prefrontal cortex tissue. We found much greater gene dysregulation within the smoking-exposed prenatal brain than the exposed adult brain. Interestingly, the significant differentially expressed prenatal genes included *GABRA4*, *NRCAM*, and *KCNN2* which have all been previously implicated in addiction (*21–24*). Moreover, genes differentially expressed in the prenatal brain and genes showing different responses to smoking exposure across the lifespan were enriched for autism spectrum disorder (ASD) genes and related intellectual impairment genes. Neuropsychiatric disorders associated with maternal smoking have extensive comorbidity with substance abuse and the differential expression associated with maternal smoking may partly explain the molecular biology that underlies the comorbidity (*25*). These results highlight dysregulation of the developing prenatal brain’s transcriptome associated with *in utero* smoking exposure and putative mechanisms of risk for neurodevelopmental disorders later in life.

## Results

We used polyA+ RNA-seq data from 207 adult, non-psychiatric and 33 prenatal postmortem brain samples with cotinine (a nicotine metabolite) or nicotine toxicology test results available to investigate transcriptomic changes associated with cigarette smoking exposure (Supplemental Table S1). Nicotine or cotinine detectability (>5 ng/mL nicotine or >10 ng/mL cotinine) was closely coupled to a case records and/or next-of-kin reported history of smoking in the proband: 87% of adult controls with nicotine/cotinine detected also had a history of smoking reported (7 samples had positive toxicology without a reported history of smoking). We did not observe significant differences in cotinine concentrations between adult active smokers with a positive next of kin history of smoking versus active smokers without a positive history (mean_+history_=279 ng/mL, N_+history_=39; mean_−history_=208 ng/mL, N_-history_=7; p=0.444) or nicotine concentrations in blood (mean_+history_=41.0 ng/mL, N_+history_=32; mean_−history_=39.8 ng/mL, N_−history_=6; p=0.936). These results suggest that the cotinine and nicotine biomarkers successfully capture the smoking-exposed phenotype.

Cigarette smoking in adults was weakly associated with increased RNA Integrity Number (RIN; p=0.052) and brain bank (p=0.024), but in the prenatal cohort smoking exposure was not associated with either RIN (p=0.76) or brain bank (p=1, Supplemental Table S1). No differences were observed at p<0.05 between the adult smoking and non-smoking groups with regards to age, ancestry, sex, or other measures of RNA quality (Supplemental Fig. S1A). In the prenatal cohort, there were trends for smoking exposure to be associated with greater gestational age (p=0.084) and higher mitochondrial mapping rate (p=0.063) (Supplemental Fig. S1B). These analyses suggest RNA quality is mildly associated with cigarette smoke exposure, a confound we sought to mitigate by adjusting for explicit measures of RNA quality and constructed surrogate variables (SVs). After filtering out lowly expressed features, 18,067 genes, 189,292 exons, 137,016 exon-exon splice junctions, and 275,885 expressed regions (ERs) (*26*) were tested for differential expression, adjusting for age, sex, ancestry, RNA quality, and latent confounds in the pre- and post-natal cohorts (see methods for details).

### Gene expression changes associated with smoking exposure in the developing brain

We performed differential expression analysis of genes and their transcript features comparing dorsolateral prefrontal cortex (DLPFC) tissue from 16 smoking-exposed compared to 17 smoking-unexposed prenatal samples. After correcting for multiple testing (FDR<10%), 14 prenatal genes had evidence of differential expression (DE), mostly at the gene level (Table 1, median absolute log_2_ fold change [LFC] = 0.618). Some of these differentially expressed genes are potentially related to nicotine dependence or addiction more broadly: *KCNN2*, *GABRA4*, and *NRCAM* (Figure 1A–C). The most significantly differentially expressed transcript feature at the gene level was *KCNN2* which encodes a small-conductance Ca^2^+-activated K+ channel (Figure 1A, p=2.60×10^−6^, LFC = −0.694). *GABRA4*, a gene coding for the alpha-4 subunit of the GABA receptor, was also differentially expressed (Figure 1B, p=1.86×10^−5^, LFC = 1.06). *NRCAM*, a gene encoding neuronal cell adhesion molecule, was differentially expressed at the gene level (Figure 1C, p=7.17×10^−5^, LFC = −0.568) and six previously annotated *NRCAM* exon-exon splice junctions were also differentially expressed (p_min_=2. 29×10, LFC_p-min_= −0.689, Supplemental Table S2). Thus, prenatal smoking exposure is associated with changes in the expression of some genes previously associated with addiction in the literature.

**Fig. 1.**
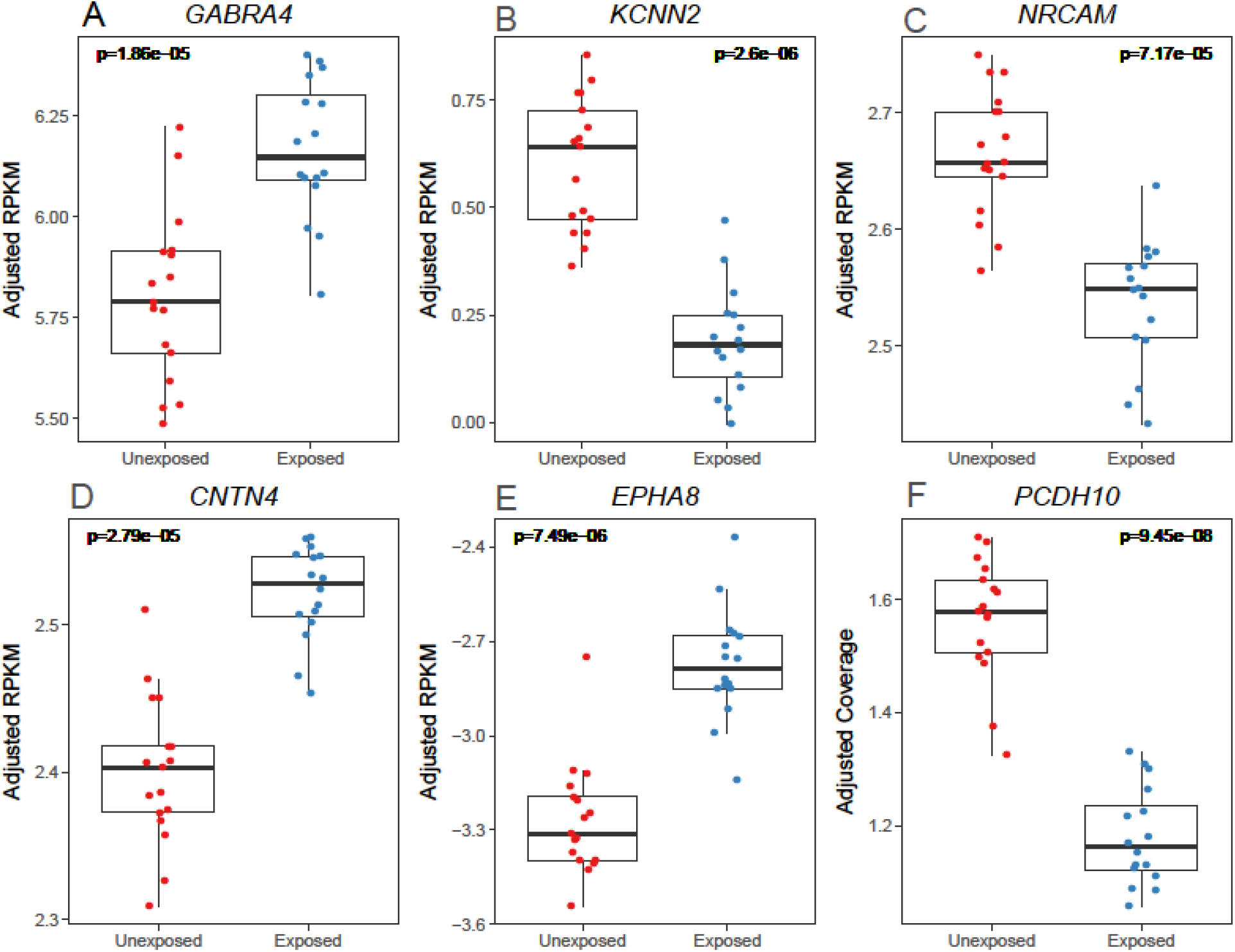
Representative differentially expressed prenatal genes. [RPKM: reads per kilobase per million]. Normalized expression levels for six genes with significant (FDR<10%) differential expression are shown for smoking unexposed and exposed prenatal prefrontal cortex samples. These representative genes have been previously implicated in neurodevelopment and neuropsychiatric disease.

**Table 1.**
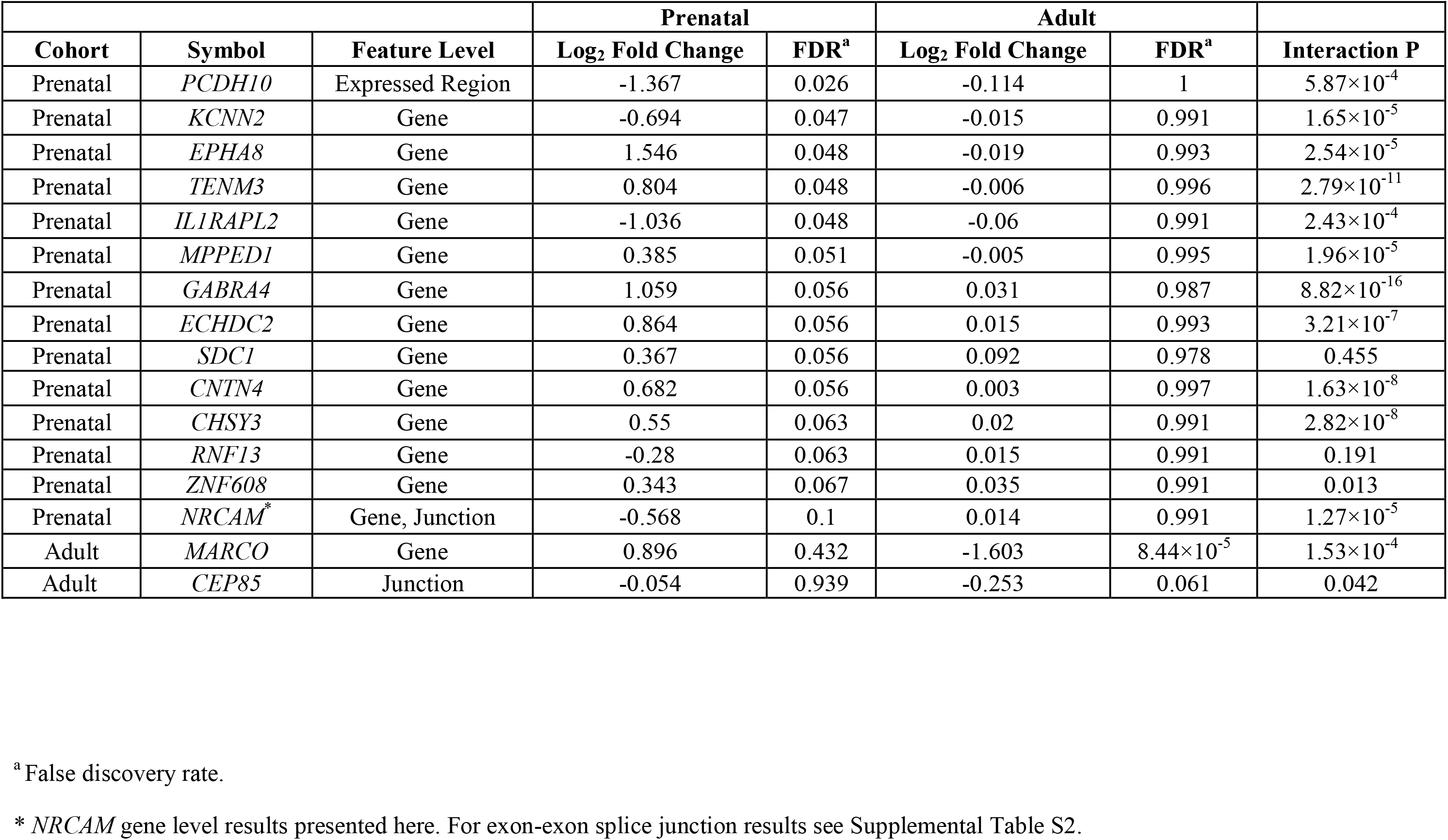
Significant differentially expressed genes between smoke unexposed and exposed human brain cortices in prenatal and postnatal cohorts (FDR<10%).

Other differentially expressed genes between the exposed and unexposed prenatal groups play a role in neurodevelopment and may be linked to neuropsychiatric disorders: *CNTN4*, *EPHA8*, and *PCDH10*(Figure 1D–F). Contactin 4 (*CNTN4*, Figure 1D, p=2.79×10^−5^, LFC=0.682) is a cell adhesion molecule involved in synaptic formation (*27*). The association with *PCDH10* was in a differentially expressed region (Figure 1E, p=9.45×10^−8^, LFC = −1.37, chr4:134,116,962-134,116,976), a 15 base pair strictly intronic region. Protocadherin-10 (*PCDH10*) encodes a cell adhesion molecule involved in synaptic elimination (*28*). EPH Receptor A8 (*EPHA8*, Figure 1F, p=7.49×10^−6^, LFC=1.55) encodes a member of the Eph family of tyrosine protein kinase receptors, which has a functional role in axonal pathfinding during neurodevelopment (*29*). A gene-level, subgroup analysis of the significant differentially expressed genes which used only African-American prenatal samples (N=30) generated effect sizes consistent with those from the original model which included 3 Caucasian smoking-exposed prenatal samples (*r*=0.99, p=1.18×10^−10^, Supplemental Fig. S2). These findings suggest, perhaps not surprisingly, that genes involved in neurodevelopment may be disrupted by prenatal exposure to cigarette smoke, a putative neurodevelopmental insult.

### More subtle gene expression changes associated with smoking exposure in the adult brain

In the adult sample, only two genes were differentially expressed between active smoker (N=150) and non-smoker groups (N=57; Table 1). *MARCO*, an immune gene, was the most significantly differentially expressed gene among adults (Figure 2A; p=4.67×10^−9^, LFC = − 0.568). In an independent cohort of DLPFC tissue from 107 patients with schizophrenia (65 active smokers, 42 non-smokers), *MARCO* was similarly differentially expressed (p=3.28×10^−5^, LFC = −1.34, Figure 2B). To assess the robustness of our model, we then stratified adult smokers into three categories: non-smokers (N=150), light-smokers (N=23), and heavy smokers (N=23), based on cotinine levels (200 ng/mL heavy-light threshold). Under this ordinal sensitivity model, *MARCO* remained significantly differentially expressed (p=6.16× 10^−10^ LFC=-1.211, Figure 2C), and our global results were highly correlated across all transcript feature levels confirming the robustness of our analysis (Supplemental Fig. S3). An exon-exon splice junction mapping to four transcripts of *CEP85* (chr1:26,595,127-26,595,950) was also differentially expressed between adult smokers and non-smokers (p=4.46×10^−7^, LFC = −0.25), but this association did not replicate in our independent schizophrenia cohort (p=0.47; Supplemental Fig. S4). Together, these findings indicate cigarette smoke exposure has subtler effects on the adult prefrontal cortex’s transcriptome than on the developing brain, despite having the larger sample size compared with the fetal sample.

**Fig. 2.**
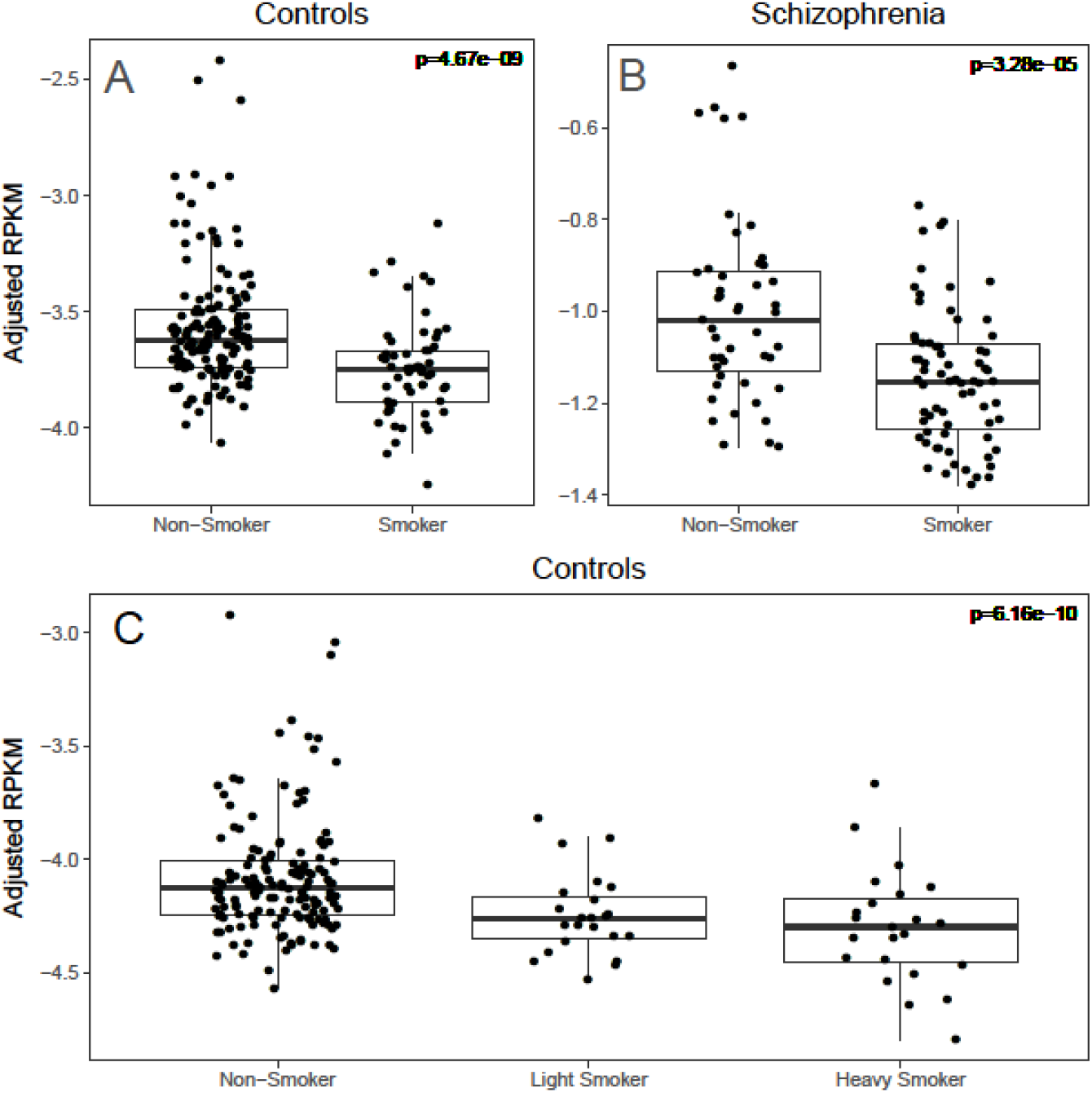
**A.** *MARCO* expression in original model across development. [RPKM: reads per kilobase per million]. *MARCO* (Macrophage Receptor With Collagenous Structure) expression was reduced in adult smokers compared to non-smokers. **B.** *MARCO* expression in the schizophrenia replication sample. *MARCO* expression was significantly reduced in smokers compared to non-smokers in a separate sample of 107 schizophrenics collected, processed, analyzed with the same pipeline as the adult controls. **C.** *MARCO* expression in sensitivity model in adults. *MARCO* expression was inversely proportional to smoking intensity under our sensitivity ordinal model in which we stratified smokers into light (N=23) and heavy (N=23) smokers.

### Significant changes in susceptibility to smoking exposure across development

After observing non-overlapping genes within the differentially expressed transcript features across the adult and prenatal cohorts associated with smoking exposure, we performed an interaction analysis to more formally determine the potentially age-dependent effects of smoking exposure in prenatal versus adult samples. While the significant main effects of smoking exposure in adulthood were limited to *MARCO* and the *CEP85* junction described above, there were 5,293 differentially expressed features corresponding to 1,315 unique Ensembl genes (Supplemental Table S3) associated with the interaction between developmental stage (pre- versus post-natal) and smoking exposure after adjusting for possible technical and biological confounds (FDR<10%). The different feature types offered both unique and convergent evidence for genes implicated as interaction genes (i.e., genes that respond differently to cigarette smoking in prenatal vs. adult cortical tissue), suggesting transcript-feature level analysis beyond the gene-level summary generates additional valuable insight into changes in the transcriptomic landscape. For instance, 87 genes were shared as significant interaction genes across all feature types, but 315 genes were solely significant for an interaction effect at the exon-level summarization (Figure 3).

**Fig. 3.**
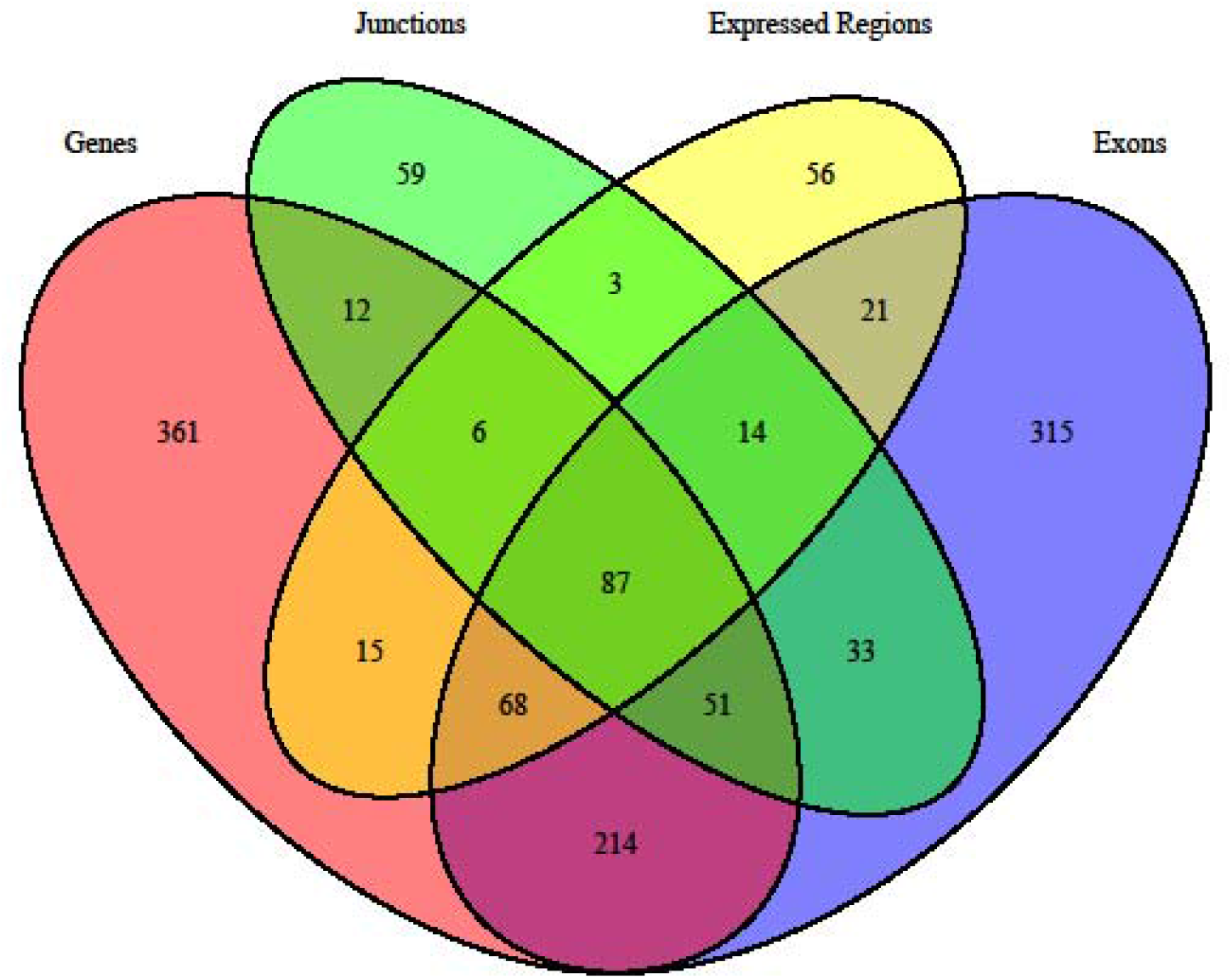
Venn diagram of differentially expressed gene overlap by feature level summarization. 87 genes are implicated across all feature level summarizations (genes, exon-exon junctions, expressed regions, and exons). Other genes are tagged uniquely by our feature level analysis such as the 56 genes that were only identified as differentially expressed regions and not by other feature-level summarizations.

We next asked whether smoking exposure’s differential effects on gene expression across the lifespan are mostly driven by significant changes in the prenatal cortical transcriptome, hypothesizing that the prenatal cortical transcriptome is more pliable than the adult one. Post-hoc analysis confirmed that more of these transcript features with a significant interaction were nominally differentially expressed (p<0.05) by smoking exposure condition in prenatal cortex (N=1,850, 35.0%) than in adult cortex (N=585, 11.1%; p=1.40×10^−8^). These findings underscore that smoking exposure exerts different effects on gene expression across the human lifespan and these effects occur more frequently in the prenatal cortex.

### Enrichment of interaction genes in diverse processes and neurodevelopmental disorders

To explore the potential functional implications of the 1,315 genes with differential effects of cigarette smoke exposure in prenatal and adult life, we performed gene ontology and pathway analyses with the sets of unique genes tagged by differentially expressed age-dependent transcript features: genes (N=752), exons (N=746), junctions (N=257), and ERs (N=285). We performed separate and convergent transcript feature level gene ontology analysis and uncovered significant enrichment of biological categories across all ontologies tested (Supplemental Table S4).

Here, “nicotine addiction” (hsa04080, p=2.04×10^−7^, q=3.15×10^−5^) was one of the most significantly enriched KEGG categories across multiple feature summarizations. Other top KEGG categories included “retrograde endocannabinoid signaling” (hsa04723, p=2.04×10^−7^, q=3.15×10^−5^) and “neuroactive ligand-receptor interaction” (hsa05033, p=2.04×10^−7^, q=3.15×10^−5^). The most significantly enriched biological processes involved “behavior” (GO:0007610, p=2.33×10^−10^, q=9.87×10^−7^), “positive regulation of nervous system development” (GO:0051962, p=1.31×10^−7^, q=2.32×10^−4^), and “synapse organization” (G0:0050808, p=2.84×10^−7^, q=2.57×10^−4^). Interestingly, using disease ontology (DO) enrichment analyses, where pre-defined gene sets are based on prior disease associations in the literature, rather than biological function/pathways, the top enriched disease ontologies (DO) were related to autism spectrum disorder (ASD), including gene sets “autism spectrum disorder” (DOID:0060041, p=8.45×10^−8^, q=2.57 ×10’^5^), “autistic disorder” (DOID:12849, p=8.45×10^−8^, q=2.57 ×10^−5^), and “pervasive developmental disorder” (DOID:0060040, p=1.25×10^−7^, q=2.57 ×10^−5^).

After observing the DO enrichment related to autism and relevant molecular processes, we performed targeted enrichment analyses using publicly available autism spectrum disorder (ASD) gene databases (*30*) and previously constructed neurodevelopmental gene sets (31) to better determine sensitivity and specificity of the ASD gene set enrichments. In prenatal samples, this ASD enrichment was driven by *IL1RAPL2, GABRA4, CNTN4, NRCAM*, and *PCDH10*being differentially expressed (Table 1, N=5/14 SFARI genes, p=3.79×10^−4^, Supplemental Table S5A). We further observed enrichment of significant age-dependent interaction genes across multiple autism gene sets including SFARI Gene (p=7.20×10^−8^), AutDb (p=1.17×10^−7^), and the curated non-syndromic ASD gene database set (p=3.77×10^−4^, Supplemental Table S5B). Interaction genes were also significantly enriched for schizophrenia genes implicated from single nucleotide variant (SNV) studies (p=0.00136) as well as nominally significantly enriched for intellectual disability genes (p=0.0115) and syndromic neurodevelopmental disorder (NDD) genes (p=0.0175; Supplemental Table S5B). The enriched NDD (N=6) were a subset of the enriched ASD genes, but the enriched schizophrenia SNV and intellectual disability genes were mostly distinct (Supplemental Fig. S5). Therefore, genes affected differently by cigarette smoke exposure across the lifespan appear enriched in autism spectrum disorder and other neuropsychiatric disorders gene sets.

### Smoking exposure’s transcriptomic pattern is similar to post-mortem brain ASD case-control differences

Next, to more fully interrogate these genes in ASD, we performed gene set enrichment analysis in post-mortem human brain tissue from a public database of patients with ASD compared to unaffected controls (*32*) by comparing the gene expression differences from our significant differentially expressed prenatal and interaction genes against ASD-related gene expression differences (Supplemental Table S6, Supplemental Fig. S6A). Our differentially expressed prenatal genes (N=14) were more highly ranked relative to other genes in terms of the ASD case-control differences in frontal cortex (BA9; p=1.35×10^−6^) and temporal cortex (BA41-42-22; p=6.49×10^−5^), but not vermis (p=0.646, Supplemental Fig. S6B). Likewise, the interaction genes tested in this ASD data (N=1,293) also showed significant enrichment for ASD case-control differences in frontal cortex (p=3.45×10^−5^) and temporal cortex (p=2.77×10^−9^), but not vermis (p=0.963, Supplemental Fig. S6C). We did not observe significant enrichment for schizophrenia case-control differences (*33, 34*) with either the differentially expressed prenatal genes (p=0.14) or the interaction genes (p=0.23). These findings suggest molecular changes in the developing human cortex associated with maternal smoking during pregnancy are analogous, at the transcriptomic scale, to the cortical pathology of ASD and these differences are, at least, partially selective.

### Smoking-associated genes form protein-protein interaction networks

We investigated the properties of protein-protein interaction (PPI) networks (*33*) constructed from proteins encoded by genes differentially expressed in prenatal development and by genes exhibiting an age-dependent response to smoking exposure. Differentially expressed prenatal genes (N=14) and the 400 most significant interaction genes mapping to unique STRING identifiers (N=384) had more PPIs than anticipated (p<0.003), and the top interaction genes also had significant subnetworks (p<2.2×10^−16^), Supplemental Fig. S7A–S7C). Next, we explored subgroups of the interaction genes which were significantly enriched for neuropsychiatric disorders. Interaction genes enriched in the SFARI autism gene database encoded proteins (N=101) had significantly more PPIs than expected (p<2.2×10^−16^), Supplemental Fig. S7D), as did genes enriched for intellectual disability (N=13, p=3.84×10^−7^, Supplemental Fig. S7E). However, interaction genes enriched for NDD had only slightly more PPI (N=6, p=0.0104, Supplemental Fig. S7F), and those enriched for in the schizophrenia SNV gene set were not enriched for PPI (N=26, p=0.532, Supplemental Fig. S7G). Further clustering of interaction genes which were linked to ASD (via SFARI Gene) highlighted significant (p<2.2×10^−16^) PPI subnetworks, such as one involving subunits of glutamate receptors, GABAergic receptors, and Ca^2+^ -voltage gated channels (Supplemental Fig. S7H). These results suggest that various groups of genes are affected by smoking-exposure, principally *in utero*, and some of these groups that are enriched for neuropsychiatric disease form functional protein association networks.

## Discussion

This study is the first, to our knowledge, to perform an extensive interrogation of the effect of smoking exposure on the prenatal cortical transcriptome. The prenatal brain is more sensitive to many environmental agents in both human and animal studies than the adult brain (*34–36*) and early neurodevelopmental exposure can associate with risk for neuropsychiatric disorders like autism, intellectual disability and schizophrenia later in life (37). Therefore, one might expect signatures of smoking exposure would be more pronounced in the developing human brain. Consistent with this hypothesis, we found 14 genes differentially expressed in the prenatal cortex compared to only 2 genes in the adult cortex.

The transcriptomic effects of smoking exposure were more noticeable in the prenatal brain despite the smaller sample size available, underscoring the effects of maternal smoking on the prenatal human brain’s sensitive developmental trajectory. Prenatal exposure to smoking was associated with changes in gene expression relevant to neurodevelopment and addiction which some studies have linked to maternal smoking during pregnancy (*38–40*). *NRCAM* encodes a neuronal cell adhesion molecule, and expression was reduced within smoking-exposed prenatal brain. This difference may have downstream consequences, as neuronal cell adhesion molecules are involved in neurodevelopment (*41–43*) and have been linked to autism (*44–46*) and schizophrenia (*47–49*). *KCNN2* levels were also reduced in the smoking-exposed prenatal group. *KCNN2* encodes small conductance Ca^2+^-activated K+ channels, and is part of a family of proteins that has been linked to substance abuse (*23, 50*).

One potential mechanism for smoking’s neurodevelopmental effects is through mediating changes to the GABAergic signaling pathway: endogenous nicotinic acetylcholine receptor signaling plays a crucial role in neuronal development by driving the GABAergic shift from an excitatory to an inhibitory neurotransmitter (*51*). Interestingly, *GABRA4*, a gene coding for the alpha-4 subunit of the GABA receptor, was previously associated with nicotine dependence in targeted studies (*21, 52, 53*). We also found nominally significant differences between the prenatal brains by smoke exposure within key genes (*CHRNA7*, *SLC12A5*, and *SLC12A2*) mediating the GABAergic transition from excitatory to inhibitory. Moreover, several other genes involved in GABAergic signaling exhibited a different response to smoking exposure across development, some of which have been previously implicated in ASD. It is plausible, although highly speculative, that the presence of nicotine due to maternal smoking during pregnancy interferes with nicotinic cholinergic-GABAergic neurodevelopmental dynamics, thus generating future neuropsychiatric risk.

Smoking exposure was also associated with changes in prenatal expression of multiple genes important in neurodevelopment such as *CNTN4* and *EPHA8*. Differentially expressed genes in the prenatal cohort were enriched in publicly available autism databases, and gene ontology analysis of the significant interaction genes (i.e., genes that were differentially expressed by smoking exposure in prenatal but not adult donors and vice versa) highlighted gene sets relevant to neurodevelopment such as synapse organization. Maternal smoking during pregnancy is associated with multiple adverse health outcomes for the offspring later in life (*37, 54-57*). These adverse outcomes include greater risk for substance abuse in adolescence which may be partially mediated via changes to the epigenetic landscape (*58*). Our findings add orthogonal molecular support to the hypothesis that maternal smoking during pregnancy may be a risk factor for neuropsychiatric disease later in life, especially for ASD.

Regardless of age, differentially expressed genes between the smoking exposed and unexposed donors could also relate to other hazardous chemicals besides nicotine, indirect effects of cigarette smoke exposure, and residual confounding by unmeasured factors associated with smoking. For example, cigarette smoke contains over 500 distinct chemicals of which 98 have been deemed dangerous to human health (*59*) and the molecular correlates of these compounds in human brain range in characterization. The genes identified in this study as differentially expressed by smoking exposure include ones that predispose to cigarette smoking susceptibility and ones that are altered by smoking or other related exposures. Nonetheless, these differentially expressed genes highlight biologically important genes and pathways that represent smoking-related changes in the brain.

Our results in DLPFC may not translate to the effects of smoking in other brain regions, as the effects of cigarette smoke exposure on the developing human cortex may vary across brain regions which have distinct cell type compositions and unique molecular profiles (*60*). Moreover, our samples are mostly of first trimester fetal brain, and the possible effects of smoking on later prenatal development may differ. Future studies should seek to determine the molecular signature of cigarette smoke exposure across multiple developing brain regions and later time periods and refine the transcriptomic footprint of exposure in the developing human cortex. Finally, the temporality and relative contributions of genes and the environment, particularly the intrauterine environment (*61*), on the observed molecular differences deserves further attention. Untangling the extent by which maternal smoking during pregnancy relates to a greater genetic burden for nicotine dependence subsequently inherited by the neonate compared to direct environmental or epigenetic insults warrants further portioning. Here, comprehensive expression quantitative trait loci mapping may help disentangle the functional consequences of predisposing genetic risk for nicotine dependence from the environmental effect of cigarette smoke exposure.

In summary, we performed transcriptome-wide scans to search for disturbances in gene expression associated with *in utero* smoking exposure. The effect of smoking exposure on gene expression was much more prominent in the developing prenatal brain than the mature adult brain. Disrupted neurodevelopment due to early environmental insults confers risk for neuropsychiatric disorders later in life, and indeed maternal smoking during pregnancy has been suggested as a risk factor for neuropsychiatric disorders in affected offspring (*3, 4*). Our study provides molecular evidence that maternal smoking during pregnancy may disrupt pathways relevant in neurodevelopment, which may influence risk of neuropsychiatric or other disease outcomes in the offspring. By characterizing transcriptomic effects of maternal smoking upon the human prenatal cortex, we have added evidence for maternal smoking as a risk factor for neuropsychiatric disease. Overall, these findings offer new molecular clues into how *in utero* smoking exposure can disrupt human cortex development and increase risk for the development of neuropsychiatric disorders in the offspring.

## Materials and Methods

### Defining smoking exposure

Presence or absence of nicotine and cotinine was determined by LC/MS-MS measurement in blood and/or cerebellar brain tissue for postnatal samples by National Medical Services (www.nmslabs.com). For prenatal samples, nicotine, cotinine, and OH-cotinine were measured by LC/MS-MS in cerebellar brain tissue as previously described (*62*). We restricted the adult cohort to non-psychiatric control samples over the age of 16 at time of death with toxicology information (N=207). “Non-smokers” (N=150) lacked detectable levels of nicotine (<5 ng/mL) and cotinine (<10 ng/mL) in blood and/or cerebellum, and did not have a history of current smoking as reported by next of kin. “Smokers” had detectable levels (>5 ng/mL nicotine or >10 ng/mL cotinine; Supplemental Fig. S8A–B) of nicotine or cotinine in blood or brain (N=57). The prenatal cohort was limited to samples before birth with toxicology information (N=33). Within the prenatal cohort, we defined “smoking-exposed” (N=16) based on having either a self-reported maternal history of sustained smoking during pregnancy or detectable levels of cotinine in brain, and “smoking-unexposed” (N=17) based on having no maternal smoking history and no detectable levels of nicotine or cotinine.

### RNA sequencing data

We used samples described in Jaffe el al. (*63*), which, briefly, were aligned to the genome and transcriptome using TopHat2 (*64*) and quantified to Ensembl v75. We used four convergent measurements of expression (“feature summarizations”) (*65, 66*): (1) gene and (2) exon counts that rely on existing gene annotation, and two annotation-agnostic approaches calculated from the read alignments – (3) read coverage supporting exon-exon splice junctions (e.g. coordinates of potentially intronic sequence that are spliced out of mature transcripts captured by a single read) and (4) read coverage overlapping each base in each sample which we have summarized into contiguous “expressed regions” (ERs) (*26*).

### Quality control and data processing

To identify and characterize potential confounders and relationships between cigarette smoking and other variables, we tested the associations between clinical and RNA quality metrics using descriptive statistics, t-tests, Fisher’s exact test, and ANOVA. Genes with mean RPKM of less than 0.1 were considered negligibly expressed and were filtered out, the gene RPKM was then transformed (log_2_(RPKM)+1) and used for principal components analysis. The association of the top five expression principle components (PCs) with clinical variables and RNA quality measures was assessed using Pearson correlations and t-tests. Samples were sequentially plotted in the first five PC spaces and these plots were inspected for the presence of outliers (0 outliers, Supplemental Fig. S9A–B). Multidimensional scaling (MDS) was performed on autosomal LD-independent genetic variants to construct genomic ancestry components on each sample, which can be interpreted as quantitative levels of ancestry – the first component separated the Caucasian and African American samples (for further details on MDS construction see Jaffe et al.) (*67*). Outlier samples (3 adult non-smokers; 2 prenatal smoking-exposed) on the first five ancestry components were dropped (Supplemental Fig. S10A–B). We calculated surrogate variables (SVs) using the sva Bioconductor package (*68*) and included them as covariates to adjust for latent confounds.

We retained genes, exons, and junctions with 10% or more samples having at least 1 count per million (cpm) that were at least partially annotated. Expressed regions (ERs; N=275,885) were defined by contiguous sequences with mean library-size normalized coverage > 5 reads per 80 million (*26*).

### Differential gene expression analyses

All features were tested under an empirical Bayes framework for significant differential expression between smoker and non-smoker adult samples, or between smoking-exposed and unexposed prenatal samples, using the moderated t-test implemented in the limma Bioconductor package (*69*). Tested features with a false discovery rate (FDR) below 10%, by feature type, were considered differentially expressed (DE).

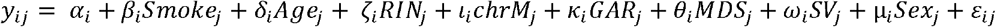

for feature *i* and donor *j* and binary smoking status *Smoke_j_*, adjusting for age, RNA Integrity Number (RIN), the proportion of reads aligning to chrM (a proxy for RNA quality), gene assignment rate (GAR, a proxy for RNA quality), the first MDS component (ancestry), surrogate variables (SVs), and the sex for each donor.

### Sensitivity analysis for smoker definitions

A sensitivity analysis was performed in our adult cohort to ascertain the robustness of results to the definition of smoker. Adult smokers with blood-based cotinine information (N=46) were stratified into light smokers (cotinine blood concentration < 200 ng/mL; N=23) and heavy smokers (cotinine blood concentration ≥ 200 ng/mL; N=23) with the threshold based on evidence from the literature (*70–72*) and also being near the median cotinine blood concentration among our samples (median concentration = 205ng/mL, Supplemental Fig. S8A). Non-smokers (N=150) were defined as discussed previously, i.e. no next-of-kin reported history and no nicotine or cotinine detected. Differential gene expression analyses were performed as described above. An ordinal model (non-smoker < light smoker < heavy smoker) was tested to determine whether differentially expressed genes had a dose-response relationship.

### Independent cohort replication

Post-mortem DLPFC brain samples from adult patients diagnosed with schizophrenia (N = 107, of which 65 were defined as smokers) obtained from the same brain repository, RNA sequenced, preprocessed, and modelled akin to the adult discovery cohort were used as a replication sample. Transcript features, tested in the replication cohort, with the same fold change directionality and nominal significance (p<0.05) as those called differentially expressed (FDR<10%) in the adult non-psychiatric control discovery cohort were considered “replicated”.

### Gene ontology

All features (genes, exons, exon-exon junctions, and ERs) were mapped to Entrez IDs (18,407 unique genes) to perform gene ontology analyses. The subsequent Entrez IDs were mapped to six different gene ontologies: KEGG(73), GO-BP (*74*), GO-MF (*74*), GO-CC (*74*), DO(*75*), and Reactome (*76, 77*). This analysis was only performed on the results of the interaction model because it was the only model with enough differentially expressed genes to ensure adequate power for statistical inference. For each ontology, the hypergeometric test for gene ontology (GO) overrepresentation was implemented via the Bioconductor package clusterProfiler independently for each feature type, except “all” which was the union of genes with significant differentially expressed features (*78*). As a reference, we used expressed genes (tested for differential expression) which had corresponding Entrez gene IDs. Multiple testing of gene ontology was controlled for with FDR correction. There was not a significant difference (p=0.21) in gene length between the differentially expressed genes (mean length = 4,504bp) and the non-differentially expressed genes (mean length = 4,711bp), therefore gene length is an unlikely source of bias for this analysis.

Targeted gene set enrichment analysis was undertaken using Fisher’s exact test of the 2x2 overlap table of differentially expressed genes against all genes present in autism databases AutDB (*30*), SFARI Gene (*79*), and neuropsychiatric gene sets from Birnbaum et al. (*31*). Only expressed genes— tested for differential expression— in our sample were included in the 2x2 contingency table.

### Re-analysis of human post-mortem ASD and schizophrenia brain data

We reprocessed sequencing reads from Parikshak et al. (*32*) using the same RNA-seq pipeline as above, and summarized expressed features relative to Ensembl v75. Within each brain region (BA41-BA42-BA22, BA9, and vermis), we calculated the moderated T-statistics of autism spectrum disorder (ASD) versus control, adjusting for gene assignment rate, brain bank, RIN, age, sex, and quality surrogate variables (*80*) using limma (*69*). The *geneSetTest* function, also from limma, was used to competitively test whether sets of genes differentially expressed by smoking exposure were differentially expressed by ASD status, separately by brain region. Schizophrenia differential expression enrichment was assessed analogously as above, using t-tests identified in two recent papers using a larger set of RNA-seq data than for the smoking analyses described here (*63, 80*)

### Protein-protein interaction network analysis

Using STRING (v. 10)(*33*) via the STRINGdb Bioconductor package, gene symbols for differentially expressed, enriched genes were mapped to unique identifiers in the STRING database (STRING score>400, homo sapiens). Functional association networks were then constructed on the basis of corresponding protein-protein interaction scores. For gene sets larger than 400 genes, the maximum number accepted by STRING, the 400 most significant (smallest P-values) genes were used. The p-values at the top of each plot represent the probability of observing an equal or greater number of protein-protein interactions by chance. These p-values were computed via the *ppi_enrichment* function from the STRINGdb package, and the set of proteins present in the STRING database was used as a background for this calculation (20,457 proteins, 4,274,001 interactions). The “fastgreedy” algorithm was used to cluster groups within resulting protein-protein interaction networks.

## Supplementary Materials

### Supplemental Figures

**Supplemental Fig. S1. A.** Association between adult smoking and potential confounds.

**Supplemental Fig. S1. B.** Association between prenatal smoking exposure and potential confounds.

**Supplemental Fig. S2.** Correlation between the original model and the African-American only model effect sizes for significant prenatal gene-level differential expression.

**Supplemental Fig. S3.** Correlation between the original adult model and the ordinal sensitivity model.

**Supplemental Fig. S4. A.** Adjusted *CEP85* expression in smokers and non-smokers within adult non-psychiatric controls.

**Supplemental Fig. S4. B.** Adjusted *CEP85* expression in adult smokers and non-smokers within schizophrenics.

**Supplemental Fig. S4. C.** Adjusted *CEP85* expression across non-smokers, light smokers, and heavy smokers within adult non-psychiatric controls.

**Supplemental Fig. S5.** Venn diagram of overlap between enriched neuropsychiatric disease genes.

**Supplemental Fig. S6. A.** Prenatal smoking exposure gene boxplots side-by-side with corresponding ASD case-control gene boxplots.

**Supplemental Fig. S6. B.** Enrichment of prenatal genes for ASD-case control moderated t-statistics in temporal and prefrontal cortex.

**Supplemental Fig. S6. C.** Enrichment of interaction genes for ASD-case control moderated t-statistics temporal and prefrontal cortex.

**Supplemental Fig. S7. A.** Protein-protein interaction network of 14 significant differentially expressed prenatal genes.

**Supplemental Fig. S7. B.** Protein-protein interaction network of the most significant interaction genes.

**Supplemental Fig. S7. C.** Four subclusters from the protein-protein interaction network of the most significant interaction genes.

**Supplemental Fig. S7. D.** Protein-protein interaction network of the significant interaction genes present in SFARI Gene.

**Supplemental Fig. S7. E.** Protein-protein interaction network of the significant interaction genes curated as intellectual disability genes.

**Supplemental Fig. S7. F.** Protein-protein interaction network of the significant interaction genes curated as neurodevelopmental disorder genes.

**Supplemental Fig. S7. G.** Protein-protein interaction network of the significant interaction genes curated as schizophrenia single nucleotide variant genes.

**Supplemental Fig. S7. H.** Four subclusters from the protein-protein interaction network of significant interaction genes present in SFARI Gene.

**Supplemental Fig. S8. A.** Distribution of the cotinine and nicotine biomarkers in adult control blood.

**Supplemental Fig. S8. B.** Distribution of the cotinine and nicotine biomarkers in adult control brains.

**Supplemental Fig. S8. C.** Association between cotinine and nicotine concentrations in adult controls.

**Supplemental Fig. S9. A.** Principle component analysis of log_2_(gene RPKM) in the adult cohort.

**Supplemental Fig. S9. B.** Principle component analysis of log_2_(gene RPKM) in the prenatal cohort.

**Supplemental Fig. S10. A.** The first five genotype multidimensional scaling (MDS) dimensions for the adult non-psychiatric samples.

**Supplemental Fig. S10. B.** The first five genotype multidimensional scaling (MDS) dimensions for the prenatal samples.

### Supplemental Tables

**Supplemental Table S1.** Demographic and technical characteristics of samples used in this study.

**Supplemental Table S2.** Prenatal significant differentially expressed *NRCAM* exon-exon splice junctions.

**Supplemental Table S3.** Transcript feature summarization results for significant interaction features.

**Supplemental Table S4.** Enrichment of interaction genes across various ontologies.

**Supplemental Table S5. A.** Enrichment of differentially expressed prenatal genes in curated gene sets.

**Supplemental Table S5. B.** Enrichment of interaction genes in curated gene sets.

**Supplemental Table S6.** Autism spectrum disorder case-control statistics for differentially expressed prenatal and interaction genes.

## Author contributions

S.A.S – performed analyses and led the writing of the manuscript

L.C.T. – performed analyses and contributed to the writing of the manuscript

C.A.M, L.J.B., B.S.M., E.O.J. – contributed to the interpretation of the results and writing of the manuscript

J.H.S. – performed data generation

A.D. – performed clinical reviews and oversaw assessment of toxicology

R.T. – performed RNA extractions

T.M.H. – performed tissue dissections, contributed to the study design, interpretation of the results, and writing of the manuscript

D.R.W. – contributed to the study design, interpretation of the results, and writing of the manuscript

D.B.H. – contributed to the study design, statistical analyses, interpretation of the results, and writing of the manuscript

J.E.K., A.E.J – co-led the study, including the design, statistical analyses, interpretation, and writing of the manuscript

## Data and materials availability

Raw and processed data will be made available through Synapse.org

## References and Notes

1 W. H. Organization, “WHO report on the global tobacco epidemic, 2011: Warning about the dangers of tobacco.,” (2011).

2 in The Health Consequences of Smoking-50 Years of Progress: A Report of the Surgeon General. (Atlanta (GA), 2014).

3 A. Stathopoulou, I. N. Beratis, S. Beratis, Prenatal tobacco smoke exposure, risk of schizophrenia, and severity of positive/negative symptoms. Schizophrenia research 148, 105–110 (2013).

4 S. Niemelä, A. Sourander, H.-M. Surcel, S. Hinkka-Yli-Salomäki, I. W. McKeague, K. Cheslack-Postava, A. S. Brown, Prenatal Nicotine Exposure and Risk of Schizophrenia Among Offspring in a National Birth Cohort. American Journal of Psychiatry 173, 799–806 (2016).

5 S. C. Curtin, T. J. Matthews, Smoking Prevalence and Cessation Before and During Pregnancy: Data From the Birth Certificate, 2014. National vital statistics reports: from the Centers for Disease Control and Prevention, National Center for Health Statistics, National Vital Statistics System 65, 1–14 (2016).

6 in The Health Consequences of Smoking: A Report of the Surgeon General. (Atlanta (GA), 2004).

7 C.-Y. Liao, Y.-J. Chen, J.-F. Lee, C.-L. Lu, C.-H. Chen, Cigarettes and the developing brain: Picturing nicotine as a neuroteratogen using clinical and preclinical studies. Tzu Chi Medical Journal 24, 157–161 (2012).

8 A. Thapar, T. Fowler, F. Rice, J. Scourfield, M. van den Bree, H. Thomas, G. Harold, D. Hay, Maternal smoking during pregnancy and attention deficit hyperactivity disorder symptoms in offspring. Am J Psychiatry 160, 1985–1989 (2003).

9 Y. Nomura, D. J. Marks, J. M. Halperin, Prenatal exposure to maternal and paternal smoking on attention deficit hyperactivity disorders symptoms and diagnosis in offspring. The Journal of nervous and mental disease 198, 672–678 (2010).

10 P. Joelsson, R. Chudal, A. Talati, A. Suominen, A. S. Brown, A. Sourander, Prenatal smoking exposure and neuropsychiatric comorbidity of ADHD: a finnish nationwide population-based cohort study. BMC psychiatry 16, 306 (2016).

11 M. G. Motlagh, L. Katsovich, N. Thompson, H. Lin, Y. S. Kim, L. Scahill, P. J. Lombroso, R. A. King, B. S. Peterson, J. F. Leckman, Severe psychosocial stress and heavy cigarette smoking during pregnancy: an examination of the pre- and perinatal risk factors associated with ADHD and Tourette syndrome. European child & adolescent psychiatry 19, 755–764 (2010).

12 H. A. Browne, A. Modabbernia, J. D. Buxbaum, S. N. Hansen, D. E. Schendel, E. T. Parner, A. Reichenberg, D. E. Grice, Prenatal Maternal Smoking and Increased Risk for Tourette Syndrome and Chronic Tic Disorders. Journal of the American Academy of Child and Adolescent Psychiatry 55, 784–791 (2016).

13 S. Leivonen, R. Chudal, P. Joelsson, M. Ekblad, A. Suominen, A. S. Brown, M. Gissler, A. Voutilainen, A. Sourander, Prenatal Maternal Smoking and Tourette Syndrome: A Nationwide Register Study. Child psychiatry and human development 47, 75–82 (2016).

14 G. L. Wehby, K. Prater, A. M. McCarthy, E. E. Castilla, J. C. Murray, The Impact of Maternal Smoking during Pregnancy on Early Child Neurodevelopment. Journal of human capital 5, 207–254 (2011).

15 K. Polanska, J. Jurewicz, W. Hanke, Smoking and alcohol drinking during pregnancy as the risk factors for poor child neurodevelopment - A review of epidemiological studies. International journal of occupational medicine and environmental health 28, 419–443 (2015).

16 Y. Jung, A. M. Lee, S. A. McKee, M. R. Picciotto, Maternal smoking and autism spectrum disorder: meta-analysis with population smoking metrics as moderators. Scientific reports 7, 4315 (2017).

17 B. R. Joubert, S. E. Haberg, R. M. Nilsen, X. Wang, S. E. Vollset, S. K. Murphy, Z. Huang, C. Hoyo, O. Midttun, L. A. Cupul-Uicab, P. M. Ueland, M. C. Wu, W. Nystad, D. A. Bell, S. D. Peddada, S. J. London, 450K epigenome-wide scan identifies differential DNA methylation in newborns related to maternal smoking during pregnancy. Environmental health perspectives 120, 1425–1431 (2012).

18 B. R. Joubert, J. F. Felix, P. Yousefi, K. M. Bakulski, A. C. Just, C. Breton, S. E. Reese, C. A. Markunas, R. C. Richmond, C. J. Xu, L. K. Kupers, S. S. Oh, C. Hoyo, O. Gruzieva, C. Soderhall, L. A. Salas, N. Baiz, H. Zhang, J. Lepeule, C. Ruiz, S. Ligthart, T. Wang, J. A. Taylor, L. Duijts, G. C. Sharp, S. A. Jankipersadsing, R. M. Nilsen, A. Vaez, M. D. Fallin, D. Hu, A. A. Litonjua, B. F. Fuemmeler, K. Huen, J. Kere, I. Kull, M. C. Munthe-Kaas, U. Gehring, M. Bustamante, M. J. Saurel-Coubizolles, B. M. Quraishi, J. Ren, J. Tost, J. R. Gonzalez, M. J. Peters, S. E. Haberg, Z. Xu, J. B. van Meurs, T. R. Gaunt, M. Kerkhof, E. Corpeleijn, A. P. Feinberg, C. Eng, A. A. Baccarelli, S. E. Benjamin Neelon, A. Bradman, S. K. Merid, A. Bergstrom, Z. Herceg, H. Hernandez-Vargas, B. Brunekreef, M. Pinart, B. Heude, S. Ewart, J. Yao, N. Lemonnier, O. H. Franco, M. C. Wu, A. Hofman, W. McArdle, P. Van der Vlies, F. Falahi, M. W. Gillman, L. F. Barcellos, A. Kumar, M. Wickman, S. Guerra, M. A. Charles, J. Holloway, C. Auffray, H. W. Tiemeier, G. D. Smith, D. Postma, M. F. Hivert, B. Eskenazi, M. Vrijheid, H. Arshad, J. M. Anto, A. Dehghan, W. Karmaus, I. Annesi-Maesano, J. Sunyer, A. Ghantous, G. Pershagen, N. Holland, S. K. Murphy, D. L. DeMeo, E. G. Burchard, C. Ladd-Acosta, H. Snieder, W. Nystad, G. H. Koppelman, C. L. Relton, V. W. Jaddoe, A. Wilcox, E. Melen, S. J. London, DNA Methylation in Newborns and Maternal Smoking in Pregnancy: Genome-wide Consortium Meta-analysis. Am J Hum Genet 98, 680–696 (2016).

19 A. Kawashima, K. Koide, W. Ventura, K. Hori, S. Takenaka, D. Maruyama, R. Matsuoka, K. Ichizuka, A. Sekizawa, Effects of maternal smoking on the placental expression of genes related to angiogenesis and apoptosis during the first trimester. PloS one 9, e106140 (2014).

20 Z. Chatterton, B. J. Hartley, M. H. Seok, N. Mendelev, S. Chen, M. Milekic, G. Rosoklija, A. Stankov, I. Trencevsja-Ivanovska, K. Brennand, Y. Ge, A. J. Dwork, F. Haghighi, In utero exposure to maternal smoking is associated with DNA methylation alterations and reduced neuronal content in the developing fetal brain. Epigenetics & chromatin 10, 4 (2017).

21 A. Agrawal, M. L. Pergadia, S. Balasubramanian, S. F. Saccone, A. L. Hinrichs, N. L. Saccone, N. Breslau, E. O. Johnson, D. Hatsukami, N. G. Martin, G. W. Montgomery, A. M. Goate, J. P. Rice, L. J. Bierut, P. A. Madden, Further evidence for an association between the gamma-aminobutyric acid receptor A, subunit 4 genes on chromosome 4 and Fagerstrom Test for Nicotine Dependence. Addiction 104, 471–477 (2009).

22 H. Ishiguro, Q. R. Liu, J. P. Gong, F. S. Hall, H. Ujike, M. Morales, T. Sakurai, M. Grumet, G. R. Uhl, NrCAM in addiction vulnerability: positional cloning, drug-regulation, haplotype-specific expression, and altered drug reward in knockout mice. Neuropsychopharmacology: official publication of the American College of Neuropsychopharmacology 31, 572–584 (2006).

23 A. E. Padula, W. C. Griffin, 3rd, M. F. Lopez, S. Nimitvilai, R. Cannady, N. S. McGuier, E. J. Chesler, M. F. Miles, R. W. Williams, P. K. Randall, J. J. Woodward, H. C. Becker, P. J. Mulholland, KCNN Genes that Encode Small-Conductance Ca2+-Activated K+ Channels Influence Alcohol and Drug Addiction. Neuropsychopharmacology: official publication of the American College of Neuropsychopharmacology 40, 1928–1939 (2015).

24 S. F. Saccone, A. L. Hinrichs, N. L. Saccone, G. A. Chase, K. Konvicka, P. A. Madden, N. Breslau, E. O. Johnson, D. Hatsukami, O. Pomerleau, G. E. Swan, A. M. Goate, J. Rutter, S. Bertelsen, L. Fox, D. Fugman, N. G. Martin, G. W. Montgomery, J. C. Wang, D. G. Ballinger, J. P. Rice, L. J. Bierut, Cholinergic nicotinic receptor genes implicated in a nicotine dependence association study targeting 348 candidate genes with 3713 SNPs. Human molecular genetics 16, 36–49 (2007).

25 D. A. Regier, M. E. Farmer, D. S. Rae, B. Z. Locke, S. J. Keith, L. L. Judd, F. K. Goodwin, Comorbidity of mental disorders with alcohol and other drug abuse. Results from the Epidemiologic Catchment Area (ECA) Study. JAMA 264, 2511–2518 (1990).

26 L. Collado-Torres, A. Nellore, A. C. Frazee, C. Wilks, M. I. Love, B. Langmead, R. A. Irizarry, J. T. Leek, A. E. Jaffe, Flexible expressed region analysis for RNA-seq with derfinder. Nucleic Acids Res 45, e9 (2017).

27 C. Betancur, T. Sakurai, J. D. Buxbaum, The emerging role of synaptic cell-adhesion pathways in the pathogenesis of autism spectrum disorders. Trends in neurosciences 32, 402–412 (2009).

28 N. P. Tsai, J. R. Wilkerson, W. Guo, M. A. Maksimova, G. N. DeMartino, C. W. Cowan, K. M. Huber, Multiple autism-linked genes mediate synapse elimination via proteasomal degradation of a synaptic scaffold PSD-95. Cell 151, 1581–1594 (2012).

29 S. Park, J. Frisen, M. Barbacid, Aberrant axonal projections in mice lacking EphA8 (Eek) tyrosine protein kinase receptors. The EMBO journal 16, 3106–3114 (1997).

30 S. N. Basu, R. Kollu, S. Banerjee-Basu, AutDB: a gene reference resource for autism research. Nucleic Acids Res 37, D832–836 (2009).

31 R. Birnbaum, A. E. Jaffe, T. M. Hyde, J. E. Kleinman, D. R. Weinberger, Prenatal expression patterns of genes associated with neuropsychiatric disorders. Am J Psychiatry 171, 758–767 (2014).

32 N. N. Parikshak, V. Swarup, T. G. Belgard, M. Irimia, G. Ramaswami, M. J. Gandal, C. Hartl, V. Leppa, L. T. Ubieta, J. Huang, J. K. Lowe, B. J. Blencowe, S. Horvath, D. H. Geschwind, Genome-wide changes in lncRNA, splicing, and regional gene expression patterns in autism. Nature 540, 423–427 (2016).

33 D. Szklarczyk, A. Franceschini, S. Wyder, K. Forslund, D. Heller, J. Huerta-Cepas, M. Simonovic, A. Roth, A. Santos, K. P. Tsafou, M. Kuhn, P. Bork, L. J. Jensen, C. von Mering, STRING v10: protein-protein interaction networks, integrated over the tree of life. Nucleic Acids Res 43, D447–452 (2015).

34 G. D. Stanwood, P. Levitt, Drug exposure early in life: functional repercussions of changing neuropharmacology during sensitive periods of brain development. Current opinion in pharmacology 4, 65–71 (2004).

35 D. Rice, S. Barone, Jr., Critical periods of vulnerability for the developing nervous system: evidence from humans and animal models. Environmental health perspectives 108 Suppl 3, 511–533 (2000).

36 P. M. Rodier, Developing brain as a target of toxicity. Environmental health perspectives 103 Suppl 6, 73–76 (1995).

37 S. Niemela, A. Sourander, H. M. Surcel, S. Hinkka-Yli-Salomaki, I. W. McKeague, K. Cheslack-Postava, A. S. Brown, Prenatal Nicotine Exposure and Risk of Schizophrenia Among Offspring in a National Birth Cohort. Am J Psychiatry 173, 799–806 (2016).

38 A. Al Mamun, F. V. O’Callaghan, R. Alati, M. O’Callaghan, J. M. Najman, G. M. Williams, W. Bor, Does maternal smoking during pregnancy predict the smoking patterns of young adult offspring? A birth cohort study. Tob Control 15, 452–457 (2006).

39 D. B. Kandel, P. Wu, M. Davies, Maternal smoking during pregnancy and smoking by adolescent daughters. Am J Public Health 84, 1407–1413 (1994).

40 R. Lieb, A. Schreier, H. Pfister, H. U. Wittchen, Maternal smoking and smoking in adolescents: a prospective community study of adolescents and their mothers. Eur Addict Res 9, 120–130 (2003).

41 G. P. Demyanenko, V. Mohan, X. Zhang, L. H. Brennaman, K. E. Dharbal, T. S. Tran, P. B. Manis, P. F. Maness, Neural cell adhesion molecule NrCAM regulates Semaphorin 3F-induced dendritic spine remodeling. The Journal of neuroscience: the official journal of the Society for Neuroscience 34, 11274–11287 (2014).

42 D. Fitzli, E. T. Stoeckli, S. Kunz, K. Siribour, C. Rader, B. Kunz, S. V. Kozlov, A. Buchstaller, R. P. Lane, D. M. Suter, W. J. Dreyer, P. Sonderegger, A direct interaction of axonin-1 with NgCAM-related cell adhesion molecule (NrCAM) results in guidance, but not growth of commissural axons. The Journal of cell biology 149, 951–968 (2000).

43 T. Sakurai, The role of NrCAM in neural development and disorders--beyond a simple glue in the brain. Molecular and cellular neurosciences 49, 351–363 (2012).

44 T. Sakurai, N. Ramoz, J. G. Reichert, T. E. Corwin, L. Kryzak, C. J. Smith, J. M. Silverman, E. Hollander, J. D. Buxbaum, Association analysis of the NrCAM gene in autism and in subsets of families with severe obsessive-compulsive or self-stimulatory behaviors. Psychiatr Genet 16, 251–257 (2006).

45 S. S. Moy, R. J. Nonneman, N. B. Young, G. P. Demyanenko, P. F. Maness, Impaired sociability and cognitive function in Nrcam-null mice. Behavioural brain research 205, 123–131 (2009).

46 T. Marui, I. Funatogawa, S. Koishi, K. Yamamoto, H. Matsumoto, O. Hashimoto, E. Nanba, H. Nishida, T. Sugiyama, K. Kasai, K. Watanabe, Y. Kano, T. Sasaki, N. Kato, Association of the neuronal cell adhesion molecule (NRCAM) gene variants with autism. The international journal of neuropsychopharmacology 12, 1–10 (2009).

47 D. Barbeau, J. J. Liang, Y. Robitalille, R. Quirion, L. K. Srivastava, Decreased expression of the embryonic form of the neural cell adhesion molecule in schizophrenic brains. Proc Natl Acad Sci U S A 92, 2785–2789 (1995).

48 L. H. Brennaman, P. F. Maness, NCAM in neuropsychiatric and neurodegenerative disorders. Advances in experimental medicine and biology 663, 299–317 (2010).

49 M. Poltorak, I. Khoja, J. J. Hemperly, J. R. Williams, R. el-Mallakh, W. J. Freed, Disturbances in cell recognition molecules (N-CAM and L1 antigen) in the CSF of patients with schizophrenia. Experimental neurology 131, 266–272 (1995).

50 J. L. Cadet, C. Brannock, I. N. Krasnova, S. Jayanthi, B. Ladenheim, M. T. McCoy, D. Walther, A. Godino, M. Pirooznia, R. S. Lee, Genome-wide DNA hydroxymethylation identifies potassium channels in the nucleus accumbens as discriminators of methamphetamine addiction and abstinence. Mol Psychiatry, (2016).

51 Z. Liu, R. A. Neff, D. K. Berg, Sequential interplay of nicotinic and GABAergic signaling guides neuronal development. Science 314, 1610–1613 (2006).

52 A. Agrawal, M. L. Pergadia, S. F. Saccone, A. L. Hinrichs, C. N. Lessov-Schlaggar, N. L. Saccone, R. J. Neuman, N. Breslau, E. Johnson, D. Hatsukami, G. W. Montgomery, A. C. Heath, N. G. Martin, A. M. Goate, J. P. Rice, L. J. Bierut, P. A. Madden, Gamma-aminobutyric acid receptor genes and nicotine dependence: evidence for association from a case-control study. Addiction 103, 1027–1038 (2008).

53 W. Y. Cui, C. Seneviratne, J. Gu, M. D. Li, Genetics of GABAergic signaling in nicotine and alcohol dependence. Human genetics 131, 843–855 (2012).

54 J. Biederman, M. Martelon, K. Y. Woodworth, T. J. Spencer, S. V. Faraone, Is Maternal Smoking During Pregnancy a Risk Factor for Cigarette Smoking in Offspring? A Longitudinal Controlled Study of ADHD Children Grown Up. Journal of attention disorders, (2014).

55 M. Melchior, R. Hersi, J. van der Waerden, B. Larroque, M. J. Saurel-Cubizolles, A. Chollet, C. Galera, E. M.-C. C. S. Group, Maternal tobacco smoking in pregnancy and children’s socio-emotional development at age 5: The EDEN mother-child birth cohort study. European psychiatry: the journal of the Association of European Psychiatrists 30, 562–568 (2015).

56 A. Talati, P. J. Wickramaratne, R. Wesselhoeft, M. M. Weissman, Prenatal tobacco exposure, birthweight, and offspring psychopathology. Psychiatry research 252, 346–352 (2017).

57 B. D. Holbrook, The effects of nicotine on human fetal development. Birth defects research. Part C, Embryo today: reviews 108, 181–192 (2016).

58 C. A. Cecil, E. Walton, R. G. Smith, E. Viding, E. J. McCrory, C. L. Relton, M. Suderman, J. B. Pingault, W. McArdle, T. R. Gaunt, J. Mill, E. D. Barker, DNA methylation and substance-use risk: a prospective, genome-wide study spanning gestation to adolescence. Translational psychiatry 6, e976 (2016).

59 R. Talhout, T. Schulz, E. Florek, J. van Benthem, P. Wester, A. Opperhuizen, Hazardous compounds in tobacco smoke. International journal of environmental research and public health 8, 613–628 (2011).

60 A. E. Jaffe, Y. Gao, A. Deep-Soboslay, R. Tao, T. M. Hyde, D. R. Weinberger, J. E. Kleinman, Mapping DNA methylation across development, genotype and schizophrenia in the human frontal cortex. Nature neuroscience 19, 40–47 (2016).

61 G. Ursini, G. Punzi, Q. Chen, S. Marenco, J. Robinson, A. Porcelli, E. Hamilton, M. Mitjans, G. Maddalena, M. Begemann, J. Seidel, H. Yanamori, A. E. Jaffe, K. F. Berman, M. F. Egan, R. E. Straub, C. Colantuoni, G. Blasi, R. Hashimoto, D. Rujescu, H. Ehrenreich, A. Bertolino, D. R. Weinberger, Placental gene expression mediates the interaction between obstetrical history and genetic risk for schizophrenia. bioRxiv, (2017).

62 D. M. Shakleya, M. A. Huestis, Simultaneous quantification of nicotine, opioids, cocaine, and metabolites in human fetal postmortem brain by liquid chromatography tandem mass spectrometry. Analytical and bioanalytical chemistry 393, 1957–1965 (2009).

63 A. E. Jaffe, R. E. Straub, J. H. Shin, R. Tao, Y. Gao, L. Collado Torres, T. Kam-Thong, H. S. Xi, J. Quan, Q. Chen, C. Colantuoni, B. Ulrich, B. J. Maher, A. Deep-Soboslay, A. Cross, N. J. Brandon, J. T. Leek, T. M. Hyde, J. E. Kleinman, D. R. Weinberger, Developmental And Genetic Regulation Of The Human Cortex Transcriptome In Schizophrenia. bioRxiv, (2017).

64 D. Kim, G. Pertea, C. Trapnell, H. Pimentel, R. Kelley, S. L. Salzberg, TopHat2: accurate alignment of transcriptomes in the presence of insertions, deletions and gene fusions. Genome biology 14, R36 (2013).

65 L. Collado-Torres, A. Nellore, K. Kammers, S. E. Ellis, M. A. Taub, K. D. Hansen, A. E. Jaffe, B. Langmead, J. T. Leek, Reproducible RNA-seq analysis using recount2. Nature biotechnology 35, 319–321 (2017).

66 L. Collado-Torres, A. Nellore, A. Jaffe, recount workflow: Accessing over 70,000 human RNA-seq samples with Bioconductor [version 1; referees: 1 approved, 2 approved with reservations]. (2017), vol. 6.

67 A. L. Price, N. J. Patterson, R. M. Plenge, M. E. Weinblatt, N. A. Shadick, D. Reich, Principal components analysis corrects for stratification in genome-wide association studies. Nat Genet 38, 904–909 (2006).

68 J. T. Leek, W. E. Johnson, H. S. Parker, A. E. Jaffe, J. D. Storey, The sva package for removing batch effects and other unwanted variation in high-throughput experiments. Bioinformatics 28, 882–883 (2012).

69 M. E. Ritchie, B. Phipson, D. Wu, Y. Hu, C. W. Law, W. Shi, G. K. Smyth, limma powers differential expression analyses for RNA-sequencing and microarray studies. Nucleic Acids Res 43, e47 (2015).

70 R. J. O’Connor, G. A. Giovino, L. T. Kozlowski, S. Shiffman, A. Hyland, J. T. Bernert, R. S. Caraballo, K. M. Cummings, Changes in nicotine intake and cigarette use over time in two nationally representative cross-sectional samples of smokers. American journal of epidemiology 164, 750–759 (2006).

71 H. J. Roethig, S. Munjal, S. Feng, Q. Liang, M. Sarkar, R. A. Walk, P. E. Mendes, Population estimates for biomarkers of exposure to cigarette smoke in adult U.S. cigarette smokers. Nicotine & tobacco research: official journal of the Society for Research on Nicotine and Tobacco 11, 1216–1225 (2009).

72 S. A. Branstetter, J. E. Muscat, Time to first cigarette and serum cotinine levels in adolescent smokers: National Health and Nutrition Examination Survey, 2007-2010. Nicotine & tobacco research: official journal of the Society for Research on Nicotine and Tobacco 15, 701–707 (2013).

73 M. Kanehisa, S. Goto, KEGG: kyoto encyclopedia of genes and genomes. Nucleic Acids Res 28, 27–30 (2000).

74 C. Gene Ontology, Gene Ontology Consortium: going forward. Nucleic Acids Res 43, D1049–1056 (2015).

75 G. Yu, L. G. Wang, G. R. Yan, Q. Y. He, DOSE: an R/Bioconductor package for disease ontology semantic and enrichment analysis. Bioinformatics 31, 608–609 (2015).

76 A. Fabregat, K. Sidiropoulos, P. Garapati, M. Gillespie, K. Hausmann, R. Haw, B. Jassal, S. Jupe, F. Korninger, S. McKay, L. Matthews, B. May, M. Milacic, K. Rothfels, V. Shamovsky, M. Webber, J. Weiser, M. Williams, G. Wu, L. Stein, H. Hermjakob, P. D’Eustachio, The Reactome pathway Knowledgebase. Nucleic Acids Res 44, D481–487 (2016).

77 M. Milacic, R. Haw, K. Rothfels, G. Wu, D. Croft, H. Hermjakob, P. D’Eustachio, L. Stein, Annotating cancer variants and anti-cancer therapeutics in reactome. Cancers 4, 1180–1211 (2012).

78 G. Yu, L. G. Wang, Y. Han, Q. Y. He, clusterProfiler: an R package for comparing biological themes among gene clusters. Omics: a journal of integrative biology 16, 284–287 (2012).

79 B. S. Abrahams, D. E. Arking, D. B. Campbell, H. C. Mefford, E. M. Morrow, L. A. Weiss, I. Menashe, T. Wadkins, S. Banerjee-Basu, A. Packer, SFARI Gene 2.0: a community-driven knowledgebase for the autism spectrum disorders (ASDs). Molecular autism 4, 36 (2013).

80 A. E. Jaffe, R. Tao, A. L. Norris, M. Kealhofer, A. Nellore, J. H. Shin, D. Kim, Y. Jia, T. M. Hyde, J. E. Kleinman, R. E. Straub, J. T. Leek, D. R. Weinberger, qSVA framework for RNA quality correction in differential expression analysis. Proc Natl Acad Sci U S A 114, 7130–7135 (2017).

